# Enhanced H295R steroidogenesis assay and its predictive value for female reproductive toxicity

**DOI:** 10.1101/2025.08.22.671440

**Authors:** Nora Bouftas, Majorie van Duursen

**Author notes:** Corresponding author* Majorie van Duursen.

## Abstract

There is a high need for accepted test methods for chemicals that affect the hormonal system, also known as endocrine disruptors (EDCs). The H295R adrenal cell line is considered the gold standard for investigating chemicals that can disrupt steroidogenesis. This method is described in test guideline 456, established by the Organisation for Economic Co-operation and Development (OECD), and currently focuses only on changes in testosterone (T) and estradiol (E2). However, the culture media from H295R cells contains a wide range of steroid hormones. To validate a more comprehensive H295R assay, we tested 15 blinded test substances in H295R cells and measured changes in the levels of 15 steroid hormones, as part of a ringtrial. The results showed that changes in the levels of the measured steroid hormones were robust and reproducible. The classification as disruptors of steroidogenesis for 14 test substances was the same based on changes in T or E2 alone, as it was based on changes in multiple steroid hormones. One test substance was negative based on changes in T and E2, but also showed changes in the alternative steroidogenesis pathway and would therefore be classified as positive. However, the relevance of this finding is difficult to determine, given the limited knowledge of the biological role of the alternative steroidogenesis pathway. While expanding the number of endpoint measurements in the H295R test method, thus measuring changes in multiple steroid hormones, does not appear to change the conclusion if a substance is (not) a steroidogenic disruptor, it may provide additional information that could help explain adverse health effects resulting from disrupted steroid hormone production. To investigate this further, an extensive literature review was conducted to evaluate the predictive value of the H295R test method for effects on female reproduction. This evaluation focused primarily on bisphenol A (BPA), BPS, BPF, and the plasticizer DEHP, as these were the areas where the most data were available for both the H295R test method and effects on female reproduction in animal studies. Although the evidence for disruption of steroidogenesis in the H295R test and the occurrence of some effects in animal studies (follicular and estrous cycle disruption) was overwhelming, establishing a direct link requires a detailed analysis. This could include examining altered levels of steroid hormones in the blood and using OECD-endorsed descriptions of mechanisms leading to adverse effects (so-called Adverse Outcome Pathways, AOPs). Based on our results, expanding the H295R assay does not appear to change the classification of steroidogenic disruptors, but could yield more mechanistic information. Combined with information from computer models, other cell-based tests, and/or animal experimental data, and supported by OECD-endorsed AOPs and AOP networks, this could contribute to clearer evidence for the link between endocrine disrupting effects of chemicals and female reproductive effects within European legislation.

## Introduction

An adequate portfolio of accepted test methods is needed to carry out the required safety assessment of chemicals prior to their entry into the European market to sufficiently protect human and environmental health against the hazard and risk of chemical substances, materials and products. In particular, the lack of accepted test methods to address chemicals that target the hormone system, i.e. endocrine disrupting chemicals (EDCs), is of great concern. In the past five years, eight EU-funded projects under the umbrella of the EURION cluster (www.eurion-cluster.eu) have worked on providing new test methods to identify EDCs covering four health domains, i.e. thyroid hormone disruption, metabolic diseases, developmental neurotoxicity and female fertility. Under this umbrella, over 100 test methods have been developed^1^. Despite the successful development of many test methods by these projects, it was recognized that additional funding is urgently needed to (further) validate these methods and ascertain uptake into regulatory processes^2^. Of the EURION projects, the FREIA project (www.freiaproject.eu) focused on the effects of EDCs on female reproductive health^3^.

One of the test methods investigated in the FREIA project is the H295R steroidogenesis assay. This cell-based assay utilizes the human adrenal cell line H295R and is used for screening chemical effects on steroid hormone formation, i.e. steroidogenesis. This test method was originally validated in 2010 to screen for effects on the production of 17β-estradiol (E2) and testosterone (T), described in OECD Test Guideline (TG) 456^4^ and is commonly used in regulatory test strategies in line with the OECD Conceptual Framework for Testing and Assessment of EDCs^5^.

The H295R cells, however, produce more steroid hormones besides E2 and T and is potentially a more versatile test model than how it is currently used. An expansion of the steroid hormone profile produced by the H295R assay can potentially provide more mechanistic information on the effects of suspected EDCs. However, for acceptance of an enhanced H295R steroidogenic assay, validation is needed to assess the robustness of changes in steroid hormone formation beyond E2 and T. Moreover, an assessment is needed to link the enhanced H295R test outcomes with adverse health effects in (regulatory) tests using experimental animals or human studies. Good, structured information to substantiate this is currently lacking, especially for an expanded steroid hormone profile. This makes the H295R assay liable for discussions regarding its applicability and predictivity. Moreover, linking outcomes of the H295R assay to female-specific endpoints would allow a better assessment of female reprotoxic outcomes in regulatory safety assessment, which is urgently needed.

The objectives of present study are two-fold:

1. Perform experimental work in Phase 2 of the validation process to evaluate the performance of the enhanced H295R steroidogenesis assay;
2. Provide insight in the value of the (enhanced) H295R assay to predict female reprotoxic effects.

### Validation of the enhanced H295R steroidogenesis assay

The formation of steroid hormones starts from cholesterol (Figure 1), involves multiple enzymatic conversions and can ultimately lead to sex steroid hormones (e.g. testosterone and estradiol) or corticosteroids (e.g. corticosterone and cortisol), dependent on the tissue where steroidogenesis takes place. More recently, a backdoor steroidogenic pathway has been discovered, that diverges at the level of 17-hydroxyprogesterone and, via conversion to 17-OH-progesterone and androstenedione, leads to the formation of dihydrotestosterone (DHT) thereby by-passing testosterone ^6^.

**Figure 1.**
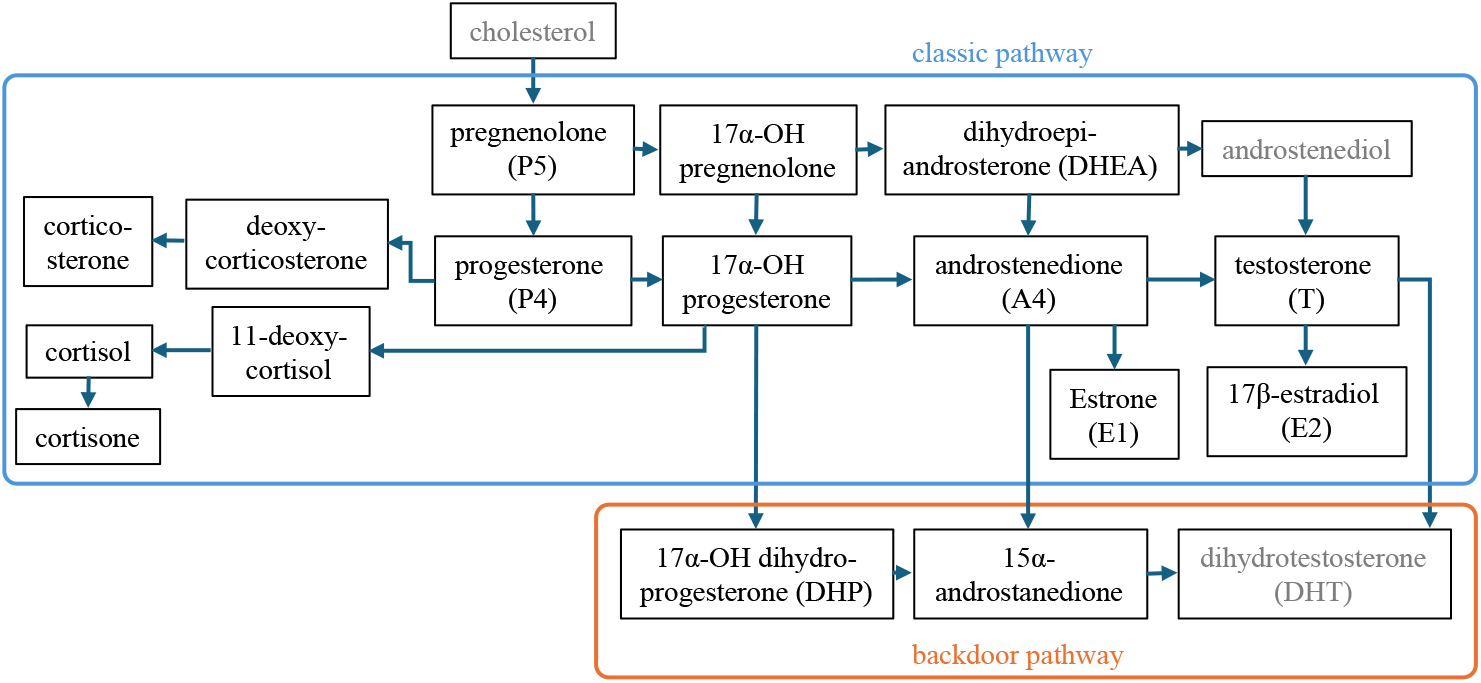
Schematic overview of the classic and backdoor steroidogenic pathways. Hormones denoted in grey were not included in the enhanced H295R steroidogenesis assay.

The adrenal-derived H295R cell line is considered a versatile model to study steroidogenesis as it can produce a broad steroidogenic profile, including some backdoor steroid hormones^7^ and does not require addition of a substrate. Studying a broader steroidome may greatly enhance the mechanistic information retrieved from the H295R assay. However, while the H295R cell line is extensively used to detect changes in T and E2 upon chemical exposure, as described in the OECD TG456, its robustness has not yet been demonstrated for other hormones.

To test the robustness of the enhanced H295R assay, a validation study was initiated to measure multiple steroid hormones using LC-MS/MS. The validation process is coordinated by PEPPER platform (https://ed-pepper.eu/en/) and supervised the by OECD Working Group of National Coordinators of the Test Guidelines Programme (WNT; project 4.159). Phase 1 of the validation has been finalized in 2024, meaning that participating labs had to demonstrate they were able to perform the assay within the established performance criteria using three reference chemicals aminoglutethimide (steroidogenic inhibitor), atrazine (steroidogenic inducer) and benomyl (negative control). November 2024, the OECD assessed the Phase 1 results and approved the continuation with the second phase of the validation. In this second phase, a ringtrial was commenced where two laboratories received 15 blind-coded test chemicals.

## Methods

Fifteen blind-coded substances were sent the Vrije Universiteit Amsterdam (VUA) by PEPPER. Test compounds were tested according to the Standard Operating Procedure (SOP), as agreed with OECD WNT and among the test laboratories. Briefly, solubility was assessed and compounds were added in a serial dilution to the H295R cells to check for cytotoxicity using an MTT assay. Where needed, the concentration range was adjusted to omit cytotoxic concentrations for steroid hormone assessment. Forskolin (1 and 10 mM) and prochloraz (0.1 and 1 mM) were used as positive controls for steroidogenic induction and inhibition, respectively. After a 48-hour exposure to the test compounds, media of the H295R cultures were collected and stored at -80°C until steroid hormone analysis. Steroid hormone levels were determined by LC-MS/MS as previously described ^8^.

## Results

### Cytotoxicity and assay performance

Of the 15 compounds tested, 5 showed cytotoxicity at the highest concentrations tested (dependent on the compound at 20 mg/ml – 200 mg/ml, data not shown). Positive controls for steroidogenic induction (forskolin) and inhibition (prochloraz) showed that the assay passed the acceptance criteria as defined in the test guideline (cell viability, limit of quantification, hormone measurement system, quality control plate, replicate measurement variability – data not shown).

### Steroidogenic disruption by the test compounds

Steroid hormone levels were determined in two experiments performed in triplicate. Of the test compounds, four compounds (Compounds 2, 12, 13, 15) caused no changes in steroid hormone levels in the canonical steroidogenic pathway. Of these negative compounds, Compound 15 caused a decrease in backdoor pathway steroid 17a-hydroxy-dihydroprogesterone. Eleven test compounds caused concentration-dependent changes in several of the hormones tested (data not shown). Notably, Compounds 3, 7 and 10 showed non-monotonic changes in hormone levels, where hormone levels initially increased and subsequently decreased. For example, Compound 3 resulted in a concentration-dependent increase in pregnenolone and 17a-OH-progesterone levels up to 200 ng/ml, but a subsequent decrease at higher concentrations (Figure 2).

**Figure 2.**
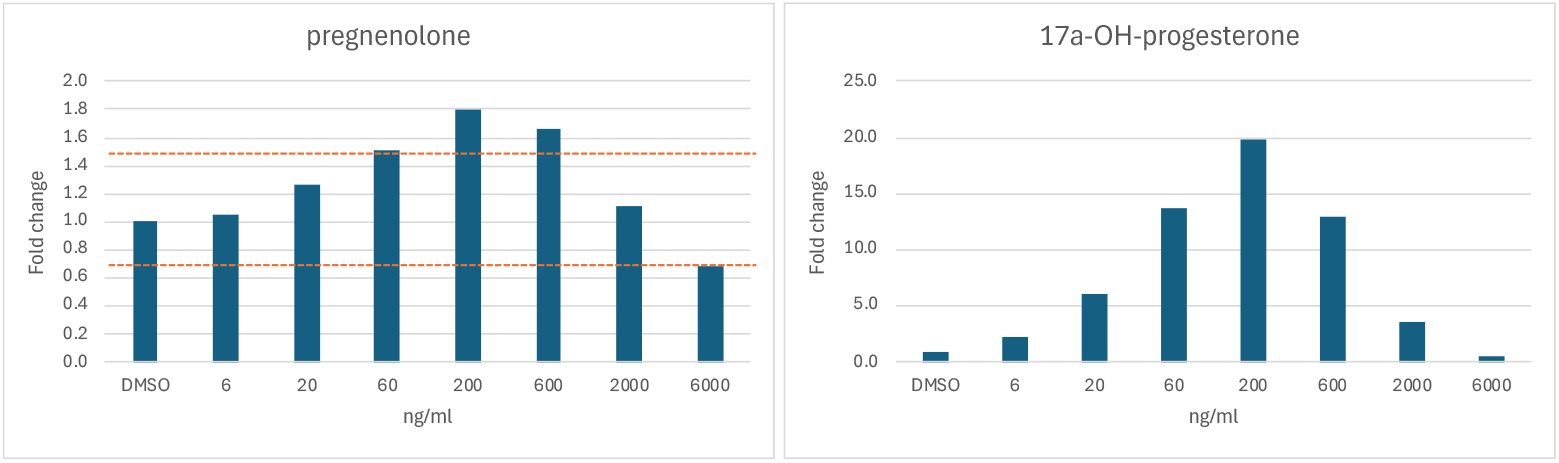
Non-monotonic changes in pregnenolone (left panel) and 17a-OH-progesterone (right panel) levels in H295R culture media upon a 48-hour exposure to Compound 3. Dashed lines indicate the OECD threshold of 1.5-fold increase or decrease in steroid hormone levels compared to vehicle (DMSO)-treated control cells.

Next, the effects of the test compounds on steroidogenesis were scored using the OECD established criteria based on changes in E2 and T levels^4^. According to the OECD Test Guideline 456, compounds score positive for steroidogenic disruption when there is a minimum of 1.5-fold change in T or E2 levels (up or down) in two consecutive concentrations. The fold-changes in hormone levels and resulting scores of the 15 test compounds are displayed in Figure 3. Based on changes in T and E2 alone, 4/15 test compounds scored *negative* and 11 compounds scored *positive* and are considered a steroidogenic disruptor. When considering the whole steroidome assessed, only the score of Compound 15 would change from *negative* into *positive* based on an increase in backdoor steroid hormone 17a-OH-DHP.

**Figure 3.**
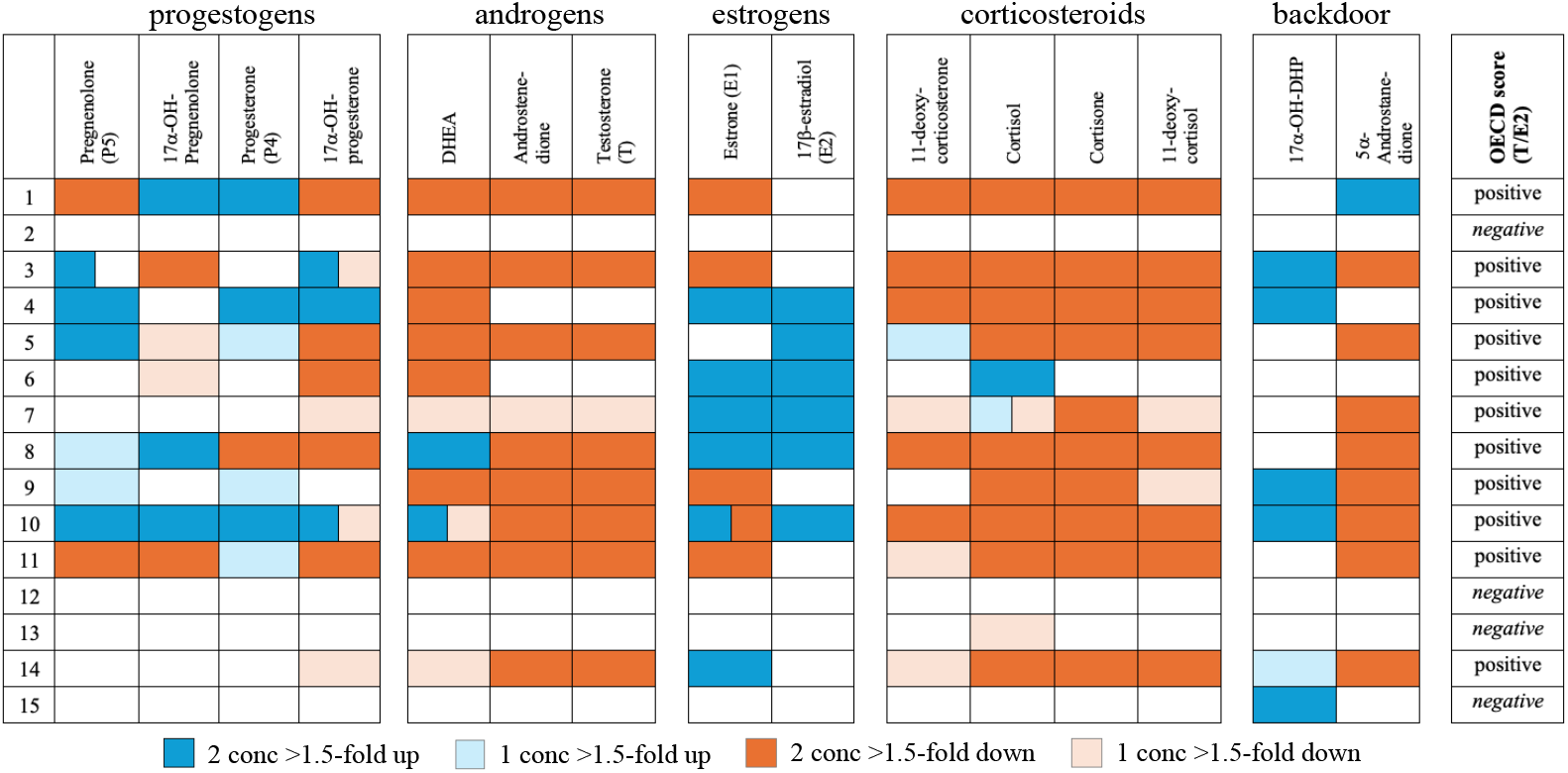
Changes in steroid hormone levels of 15 test compound in the H295R assay. Colors indicate the direction of change in hormone level (blue denotes an increase and orange a decrease) that showed more than 1.5-fold change up or down, compared to vehicle-treated controls. Dark colors indicate that 2 consecutive test concentrations exceeded the 1.5 threshold and light color indicates that 1 concentration exceeded the threshold. OECD TG456 criteria were used to score steroidogenic potential as positive or negative, based on changes in testosterone (T) and/or 17b-estradiol (E2), based on 2 independent experiments in triplicate (last column). Non-monotonic changes in hormone levels for Compounds 3, 7 and 10 are indicated as bicolored boxes.

## Discussion

The H295R assay is a versatile model to study effects of chemicals on steroidogenesis as it comprises a broad steroidogenic profile and does not require addition of a substrate. Assessing a broader steroidome, beyond currently prescribed T and E2 in the OECD TG456, was shown to be robust and reproducible. However, to further conclude on the validity of the enhanced H295R assay, the results need to be uncoded and compared with the results from the other laboratory in the ringtrial. Our results show that many changes occur early in the steroidogenic pathway, i.e. in progestogen levels, which are currently not addressed in regulatory test strategies. An enhanced H295R assay may increase the mechanistic understanding on the direction of changes in the steroidogenic pathway and where pathway effects may take place i.e. which enzymes may be affected. Notably, some compounds caused a non-monotonic response in some steroid hormone levels. Such effect has previously been described for ketoconazole, which causes an initial increase in progesterone and corticosterone levels and a subsequent decrease at higher test concentrations in the H295R assay^7,9,10^. This effect has been attributed to the differential effects ketoconazole can have on different enzymes in the steroidogenic pathway. We showed recently that H295R assay outcomes appeared to support some of the in vivo effects on circulating steroid hormones in rats perinatally exposed to ketoconazole, but only when a broad steroidomic profile and pharmacokinetics were considered ^7^. As the test compounds in this study were blind-coded to the researchers, it is not possible at this point to explain the non-monotonic responses observed. This phenomenon does require further exploration, especially in light of regulatory decision-making based on H295R outcomes.

Based on our results, expanding the current test guideline of the H295R steroidogenic assay to include more steroid hormones, particularly progestogens, corticosteroids and backdoor steroid hormones, seems relevant as it can provide information on changes in multiple steroid hormones that are associated with adverse health outcomes. However, expanding the number of steroid hormones as readouts in the H295R assay did not result in different classification of the test compounds as being a steroidogenic disruptor, except for Compound 15. It should be noted that this classification of Compound 15 changed based on changes in backdoor steroid hormone 17a-OH-DHP. The backdoor steroidogenic pathway has gained more attention in recent years for its potential role in adrenal and gonadal function and development. For example, backdoor pathway steroids were previously detected in human fetal ovarian tissue cultures after gestational week 13^11^ and have been linked with prenatal virilization of girls affected by congenital adrenal hyperplasia^12^. Despite the limited knowledge about the role of the backdoor steroidogenic pathway in development and reproduction, assessing effects on backdoor steroid hormones in the H295R assay may potentially be important for identifying chemicals with early developmental effects in the future.

### Link between altered steroidogenesis and female reproductive toxicity

Improved understanding of effects on steroidogenesis may support better identification of EDCs that cause adverse health effects. This is especially important regarding female reproductive toxicity, for which the lack of sensitive endpoints to capture EDC-mediated effects on the ovary is still of high concern^3,13^, particularly considering the increasing evidence that links exposure to EDCs with female reproductive effects^14^. At the same time, there is a push to move away from animal test methods, also in regulatory safety assessment, which inevitably places greater reliance on new approach methodologies (NAMs), including in vitro assays. Progress in NAM-based identification of EDCs may be supported by increased insights on the link between readouts from in vitro studies with adverse outcomes in vivo^15^. To this end, we evaluated the link between changes in steroidogenesis, determined by the H295R steroidogenesis assay, and female reproductive outcomes in experimental animal studies.

## Methods

### Literature search, ECHA dossiers data mining, and selection of studies

A literature search was performed using PubMed using a single concept search with the simple search string “H295R” on November 8th, 2024. Regarding the in vivo studies, the search term included the in vivo outcome and a specific compound class: “(female repro^*^ toxi^*^) AND compound”. Both searches were limited to the past 10 years. Two classes of chemicals were searched for the in vivo studies; bisphenols and phthalates. The search for bisphenols and female reproductive toxicity was done on January 8th, 2025, and the search for phthalates and female reproductive toxicity was performed on January 24^th^, 2025. Clear inclusion and exclusion criteria were defined for the H295R studies (Table 1) and in vivo female reproductive toxicity studies (Table 2). An initial screening of the abstracts was performed. If exclusion could not be clearly defined, an examination of the full text was then undertaken.

**Table 1.** Inclusion and exclusion criteria for the selection of in vitro H295R assay studies.

| Inclusion criteria | Exclusion criteria |
| --- | --- |
| Research articles | Other types of articles |
| English | Other languages |
| Endpoint -hormone measurement | Other endpoints |

**Table 2.** Inclusion and exclusion criteria for the selection of in vivo female reproductive toxicity studies.

| Inclusion criteria | Exclusion criteria |
| --- | --- |
| Research articles | Other types of articles |
| English | Other languages |
| Mammals | Other animal models |
| Female reproductive toxicity endpoints | Other endpoints |
|  | Mixtures |

ECHA dossiers for bisphenols and phthalates were also screened. The list of substances of very high concern (SVHC) was filtered to only include endocrine disrupting chemicals related to human health (Article 57(f)). Dossiers of bisphenols and phthalates were screened for H295R data and female reproductive toxicity data. Studies associated with these data were then selected for subsequent steps.

### Reporting of the information

Information from the selected, screened studies was collected into two tables: one for in vitro H295R assay studies and a table for the in vivo female reprotoxic articles (not shown). The information recorded for H295R studies includes the specific compound, the concentrations of the compound, time of exposure, the hormones measured, and the effect of the exposure including the fold change and if the effect is observed in two consecutive concentrations, following the OECD guidelines in the TG456^4^. The information of the in vivo studies was registered per compound class and information on type of species, doses of the compound used, exposure route, exposure window, number of animals used, types of endpoints and the effect were recorded.

### Assessment of the reliability

All selected studies were assessed for reliability using the web-based Science in Risk Assessment and Policy (SciRAP) tools available through the Karolinska Institute (https://ki.se/en/imm/scirap-science-in-risk-assessment-and-policy/scirap-tools). Separate SciRAP tools were used to assess the reporting and methodological qualities of the in vivo and in vitro studies, with each criterion being evaluated as Fulfilled, Partially Fulfilled, Not Fulfilled, or Not Determined. The SciRAP tools were used to obtain a score from 0 to 100 and translated into Klimisch categories for the reliability: “reliable without restriction”, “reliable with restriction”, “not reliable”, or “not assignable” as described in Suppl Table S1. Among the criteria available in the SciRAP tools, key criteria were selected as the most crucial criteria for the assessment of the reliability of the studies.

### Overall weight of evidence assessment

The weight of evidence for the H295R studies were assessed for changes in 17β-estradiol (E2), testosterone (T), progesterone (P4), and androstenedione (A4) levels. The evidence from the in vivo studies were divided by endpoint, including disrupted estrous cycle, impacted follicles, ovarian weight, and timing of vaginal opening. The lines of evidence for each endpoint were evaluated for each compound. A minimum of three studies were considered sufficient for categorization, and both positive and contradictory data were included. The description of the categories is presented in Suppl Table S2.

## Results

### Study selection in literature and ECHA dossiers

An initial simple search on PubMed for “H295R” resulted in 466 publications. After evaluation of these studies using the inclusion and exclusion criteria (Table 1), a total of 126 publications were included for further evaluation. Based on this evaluation, two main groups of chemicals were found to be the most common; Bisphenols and Phthalates. In total, 8 publications were identified that described bisphenol exposure and 7 studies on phthalate exposure using the H295R assay (Figure 4). These two classes of compounds were used to define the in vivo female reprotoxic study searches.

**Figure 4:**
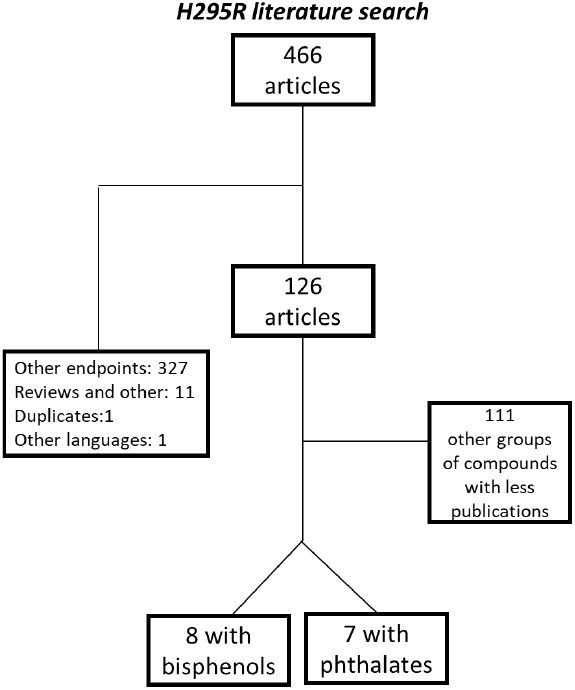
Selection process of articles in PubMed for H295R data.

As this study aimed to demonstrate a link between the H295R data and female reproductive toxicity, a search for female reproductive toxicity in relation to bisphenols and phthalates was performed. The search string for the PubMed search “(female repro^*^ toxi^*^) AND Bisphenol”, resulted in 651 publications while “(female repro^*^ toxi^*^) AND phthalate” resulted in 669 publications. Application of the inclusion and exclusion criteria (Table 2) resulted in inclusion of 36 articles for Bisphenols and 43 articles for Phthalates (Figure 5).

**Figure 5:**
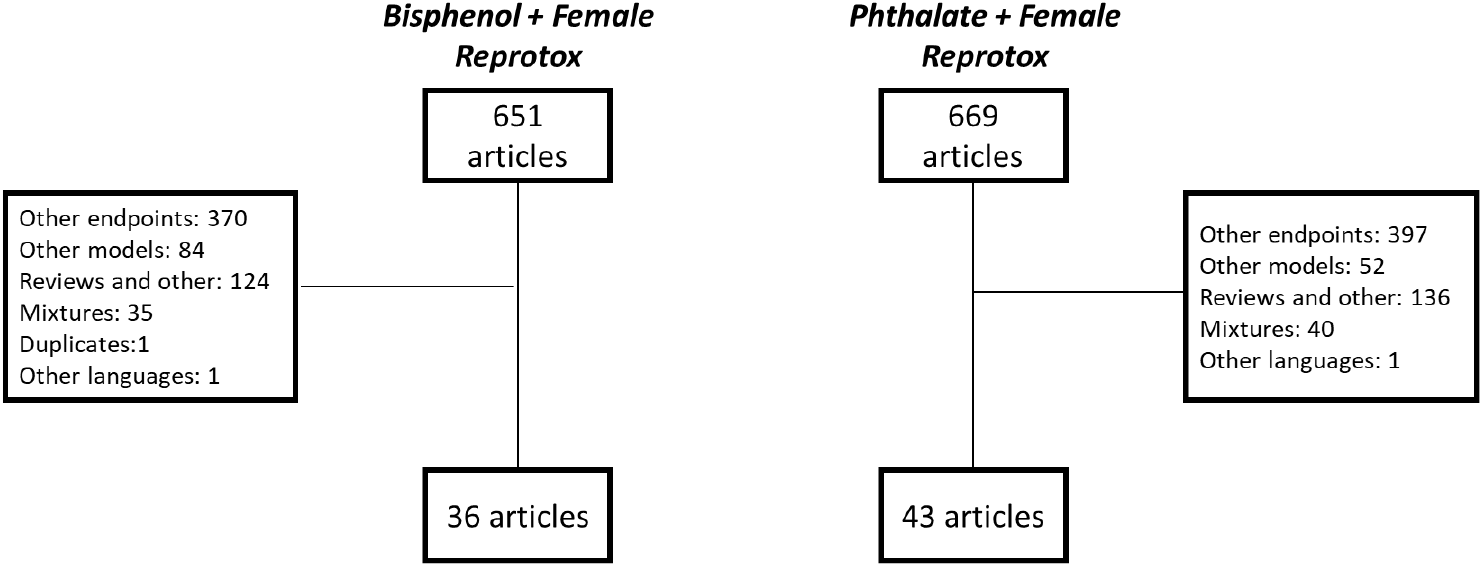
Selection process of articles in PubMed for Bisphenol or Phthalate exposure and female reproductive toxicity endpoints.

Regarding the ECHA SVHC dossiers, only the compound dossiers that included information on H295R assay results were further analyzed. Some studies described were referenced as unpublished and therefore could not be included in the present evaluation. Additionally, two H295R studies on bisphenol exposure^16,17^, and three in vivo studies also with bisphenol exposure ^18–20^ were already found in our literature search. The studies describing phthalate exposure in the ECHA dossiers were published prior to the literature search time frame of 10 years. As a result, the search through the ECHA dossiers resulted in three additional H295R publications, two relating to bisphenol exposure ^21,22^ and one phthalate exposure ^23^. For the in vivo studies, one additional publication was obtained for bisphenols ^24^, and three for phthalate exposure ^25–27^.

### Assessment of selected papers

Due to the limited number of available publications with relevant results in both in vitro and in vivo studies for other compounds, only BPA, BPS, BPF and DEHP data were analyzed in more detail.

Using the SciRAP tool, the in vitro and in vivo studies were assessed for reliability. The key criteria for the H295R studies include control, time of exposure (48h), and the measurement of cytotoxicity. To align with the OECD TG456, additional key criteria were included: a 1.5-fold change for 2 consecutive concentrations, and effect assessment in an unstimulated cell culture (i.e. effects were compared to vehicle-treated control cells). These additional key criteria were considered essential for the reliability of the effect on hormone levels although this limited the number of studies used in the overall analysis. In total for bisphenols, there were four “not reliable” ^16,28–30^ studies, 2 “reliable with restriction” ^22,31^, and two “reliable without restriction”^17,21^. Regarding DEHP exposure on H295R cells, the assessment included one “not reliable”^23^, two “reliable with restriction”^32,33^, and one “reliable without restriction”^34^.

For the reliability assessment of the in vivo studies, the key criteria included; a control group, reliable and sensitive test methods used, and a sufficient number of animals used. In total, with a focus on BPA, BPS and BPF, four “not reliable”^35–38^, twenty “reliable with restriction”^19,20,24,31,39–54^, and ten “reliable without restriction”^18,55–63^ publications were utilized in the overall weight of evidence analysis. Finally, assessment of in vivo studies with DEHP exposure resulted in ten “not reliable”^64–73^, sixteen “reliable with restriction” ^25,74–88^, and seven “reliable without restriction” ^89–95^.

### Weight of evidence analysis

#### In vitro disruption of steroidogenesis - H295R assay

The assessment of the H295R data was only performed when a positive response was observed, i.e. at least a 1.5-fold change in T or E2 levels for two consecutive concentrations, which reduced the number of publications and results included, but focused on relevant information for the present evaluation. The evidence for weak positives (i.e. two positive and one equivocal result in three experiments) was not included, while the contradictory evidence includes both negative and opposite results. The lines of evidence were then categorized as *strong, moderate*, or *weak* (Suppl Table S2), only when a minimum of 3 studies was available.

As seen in Table 3, the hormones with the most relevant data include mainly changes in 17b-estradiol and testosterone levels. Data on other hormone levels were included when available, and these included androstenedione (A4), progesterone (P4), and cortisol. The evidence that BPA exposure leads to decreased testosterone (T)^16,17,21,22,28,30^ and androstenedione (A4) levels^21,22,28^ in the H295R assay was considered to be *strong*. On the other hand, there is *weak* evidence that BPA exposure leads to increased levels of 17b-estradiol (E2) in the H295R assay^16,21,22^. Regarding BPS, there is *strong* evidence for reduced cortisol levels^16,21,28^, while no effect was found on E2 levels^16,17,21,29^. The evidence for BPS leading to decreased T levels in the H295R assay was considered *weak*, since all three studies were judged as “not reliable”^16,28,29^. For BPF, the evidence that BPF exposure leads to increased E2^17,21,28,31^ and increased P4^16,21,28^ in the H295R assay was considered *strong* for both hormones. Finally, the H295R data for DEHP exposure indicated *moderate* evidence for increased E2 due to the absence of a study judged as “reliable without restriction”^23,32,34^. The evidence for decreased T levels by DEHP was considered *very weak*, since only one study showed a decrease in T levels which was judged as ‘not reliable’^23^, while two other studies showed negative results^32,34^.

**Table 3.** Weight of evidence analysis of the H295R data. Categorization criteria are given in Table S2. The numbers in brackets indicate the number of studies the assessment was based on.

|  | Increased E2 | Decreased T | Decreased A4 | Decreased Cortisol | Increased P4 |
| --- | --- | --- | --- | --- | --- |
| **BPA** | WEAK (n=3) | STRONG (n=6) | STRONG (n=3) | - | - |
| **BPS** | No impact (n=4) | WEAK (n=3) | - | STRONG (n=3) | - |
| **BPF** | STRONG (n=4) | - | - | - | STRONG (n=3) |
| **DEHP** | MODERATE (n=3) | Very WEAK (n=3) | - | - | - |

#### Effects on female reproductive endpoints in vivo

Assessment of the lines of evidence for female reproductive toxicity was performed for the four most data-rich endpoints, namely impacted follicles, disrupted estrous cycle, early vaginal opening, and decreased ovarian weight (Table 4). A range of doses as well as different types of exposure windows and routes were included in this analysis with evidence from rats and mice, and one study with piglets.

**Table 4.** Weight of evidence analysis of the in vivo female reproductive toxicity data. Categorization criteria are given in Table S2. The numbers in brackets indicate the number of studies the assessment was based on.

|  | Impacted follicles | Disrupted estrous cycle | Early vaginal opening | Decreased ovarian weight |
| --- | --- | --- | --- | --- |
| **BPA** | STRONG (n=15) | STRONG (n=12) | MODERATE (n=9) | MODERATE (n=10) |
| **BPS** | STRONG (n=7) | STRONG (n=4) | WEAK (n=7) | MODERATE (n=4) |
| **BPF** | - | - | - | WEAK (n=3) |
| **DEHP** | STRONG (n=21) | MODERATE (n=14) | WEAK (n=9) | WEAK (n=13) |

The analysis resulted in *strong* evidence for BPA exposure leading to impacted follicles^19,20,31,35,36,38,39,44,50,52,59,60,63^ and disrupted estrous cycle^19,35,44,45,49,54,59,61,62^. There were conflicting studies found with no effect observed for the follicle population in two studies^47,53^ and no impact on estrous cyclicity in three studies^37,50,53^. However, the differences in effect could be explained by differences in exposure windows and doses. Due to the larger number of studies showing an effect on follicles and estrous cycle compared to number of conflicting studies, the evidence for these endpoints was classified as *strong*. The evidence for decreased ovarian weight^20,39,42,59,61,63^ was considered *moderate* as two studies^47,50^ found no effect on ovarian weight while one study^60^ showed an increased ovarian weight index, and one other showed a slight non-significant increase^46^. The evidence linking BPA exposure to early vaginal opening^18,19,42,43,45,46,49^ was deemed *moderate* due to two studies showing a delay in vaginal opening of which one was observed in the F3 generation ^54^ and one could not be explained ^63^.

For BPS, the evidence for a disruption of the estrous cycle in vivo was considered *strong*. Four studies^19,41,49,55^ measured a disruption of cyclicity with no conflicting data. This includes effects observed in F1 generation and one study measuring effects in the F3 generation^49^. There was *strong* evidence indicating impacted follicles^19,20,40,47,63,96^ upon exposure to BPS; with only one contradicting study showing no effects ^58^. Regarding ovarian weight, three studies showed decreased ovarian weight ^20,41,63^ and one study showed no effects ^47^, thereby leading to the overall classification as *moderate*. Early vaginal opening was observed in three studies^19,49,55^, but no effect was seen in two studies^18,58^, and one study showed delayed vaginal opening ^63^. Additionally, one of the three studies on BPS that described early vaginal opening also showed delayed vaginal opening at a lower dose^19^. Considering the amount of conflicting data compared to confirming data, the evidence for BPS causing earlier vaginal opening was categorized as *weak*.

There were very few studies identified that determined the impact of BPF on female reproductive endpoints. Of the studies included, only three could be included in the weight of evidence assessment as they examined the same endpoint, namely ovarian weight. However, the evidence was categorized as *weak* as one study showed a decrease in absolute ovarian weight but non-significant change in relative weight^56^, one study measured a clear decrease of paired ovarian weight ^20^ and one study observed no effect on ovarian weight^47^.

There is *strong* evidence suggesting that DEHP exposure leads to altered follicle development and follicle assembly^65,66,70–75,78–80,85,87–89,91,92,97,98^. Although there are two studies^76,86^ providing contradictory data for impacted follicles, it was still categorized as *strong* because of the overwhelming number of studies showing an effect and the conflicting results could be explained by a difference in animal species and exposure windows. The evidence for a disrupted estrous cycle ^64,67,69,74,78,80,87,89–93,98^ by DEHP was considered *moderate*, because three of these studies were judged as not reliable and there was one conflicting study^88^ where contradictory results could not be explained. The evidence for early vaginal opening^69,78,94^ and decreased ovarian weight^72,78,88,91,93,98^ by DEHP was considered *weak*, as there was a larger proportion of contradictory studies^25,67,71,73,74,76–78,82,84,88^ compared to studies demonstrating these two effects.

#### Changes in serum hormone levels in vivo

In addition to the in vivo endpoints assessed above, some studies also described serum hormone level measurements, mainly E2, T and P4. This data was also analyzed using the weight of evidence categorization when a minimum of three studies was available (Table 5). Only one in vivo study measured serum hormone levels upon BPF exposure, and therefore was not included in this analysis.

**Table 5.** Weight of evidence analysis for changes in serum hormone levels in females. Categorization criteria are given in Table S2. The numbers in brackets indicate the number of studies the assessment was based on.

|  | Serum E2 | Serum T | Serum P4 |
| --- | --- | --- | --- |
| BPA | Decrease and Increase:<br>WEAK (n=12) | Increase:<br>WEAK (n=7) | Decrease:<br>WEAK (n=7) |
| BPS | Increase:<br>WEAK (n=4) | Increase:<br>MODERATE (n=3) | - |
| DEHP | Decrease:<br>MODERATE (n=14) | Decrease:<br>MODERATE (n=8) | Decrease:<br>WEAK (n=9) |

The evidence for an effect of BPA on serum hormone levels was considered *weak* due to the considerable amount of contradictory data that could not be explained by differences in experimental design. Twelve studies showed changes in serum E2 levels upon BPA exposure, but these included both decreased and increased levels. Of the studies that showed a decrease in E2 serum levels, two were considered “reliable without restriction”^60,61^, one “reliable with restriction”^31^, and one “not reliable”^38^. For the studies that showed an increase in E2 serum level upon BPA exposure, one was judged as “reliable without restriction” ^18^ and three “reliable with restriction”^42,46,49^. Four of the twelve studies with BPA described no effect on serum E2 levels^19,37,44,53^. Therefore, the evidence that BPA causes changes in E2 levels was considered *weak*. Only two studies described an increase in T serum levels for BPA ^42,46^, while five observed no effect on serum T levels^18,19,38,44,49^. Additionally, four BPA studies described decreased P4 serum levels^31,37,60,61^, one study showed an increase in P4^46^, and two showed no effect^38,44^. Therefore, the evidence was categorized as *weak*.

Evidence that exposure to BPS leads to increased levels of serum T was considered *moderate*. Two studies showed an increase in serum T levels^18,19^, and one study showed no effect on serum T levels^49^. However, this difference could likely be explained as the study showing no impact focused on effects in the F3 generation. Therefore, the evidence for BPS leading to increased serum T levels was classified as *moderate*. The link between BPS exposure and increased serum E2 levels was considered *weak*, with two studies showing an increase ^18,49^, one measured a decrease in serum E2 levels^40^, and one did not observe an effect ^19^. These conflicting results could not be explained by differences in experimental design and therefore, this evidence was categorized as *weak*. No studies were found that analyzed P4 serum levels upon BPS exposure.

The evidence linking exposure to DEHP with decreased serum levels of E2 and T was considered *moderate*. Nine studies^64,65,72,75,77,78,80,86,93^ showed decreased serum E2 levels, while four studies indicated an increase in serum E2 levels^70,78,79,98^, and one study did not observe an effect ^87^. Although conflicting results were found, these could be explained by differences in species, doses, and exposure windows. The analysis for a DEHP-mediated effect on serum T levels showed four studies^70,72,93,98^ with a clear decrease in T whereas four studies^76,77,86,87^ described no effect on serum T levels. These conflicts could be explained by differences in species, exposure windows and doses, categorizing the overall evidence to be *moderate* for DEHP exposure leading to decreased serum T levels. Finally, evidence suggesting DEHP exposure leads to decreased serum P4 hormone levels was judged as *weak*. Five studies measured a decrease in P4 serum levels^64,65,72,86,93^, including three studies that were considered “not reliable”^64,65,72^. On the other hand, three studies measured an increase in P4 serum levels^76,80,87^, and one observed no effect^70^. There were no obvious differences in experimental design that could explain these conflicts, therefore categorizing the evidence that DEHP exposure leads to decreased P4 levels as being *weak*.

## Discussion

Generally, the H295R data retrieved showed *strong* evidence for steroidogenic disruption based on at least one hormone for BPA, BPS, BPF and *moderate* for DEHP. It should be noted that stringent inclusion criteria were applied, where only H295R data was included that showed at least two effect concentrations, which limited the number of studies included in the analysis. Of the hormones where evidence was *strong*, currently only changes testosterone (observed for BPA) is included in the H295R steroidogenesis test guideline^4^. Based on this, it is opportune to conclude that addition of more steroid hormones would increase the evidence base to assess the endocrine disrupting properties of a substance under evaluation. Evidence to link exposure to BPA, BPS, BPF and DEHP on female reproductive outcomes in in vivo studies was considered *weak* to *strong*, depending on the endpoint assessed.

Despite strong evidence for steroidogenic disruption from the H295R data and strong evidence for some female reproductive endpoints, a direct link between these changes is not evident without further substantiation (Figure 6). Adverse Outcome Pathways may help to further support this link. For example, our evaluation of the H295R data suggest an induction of aromatase activity, as it indicated a decrease in T (strong evidence) and increase in E2 (weak evidence) for BPA. In the AOPWiki (https://aopwiki.org), there is one AOP that describes the pathway from aromatase induction leading to estrogen receptor alpha activation via increased estradiol (<u>AOP561</u>). However, this AOP is still under development and provides no link to an adverse outcome. Still, changes in sex steroid hormone levels will affect nuclear receptor activation, e.g. of the androgen receptor (AR) and estrogen receptor (ER), and multiple AOPs are available that link AR antagonism and ER agonism to decreased fertility in mammals and fish. It should be noted that compounds can also directly act on AR/ER activation. This was not assessed in present study. Additionally, steroidogenesis in the in vitro H295R assay obviously lacks feedback regulation of the hypothalamic-pituitary-gonadal/adrenal axis, which may limit the translatability of the observed in vitro effects to the in vivo situation. Our evaluation showed that the in vitro observed changes in T and E2 levels by BPA (^–^T/-E2) are opposite to what is observed in circulating serum levels in vivo (weak evidence for -T/^–^E2). This aligns with a thorough assessment of the endocrine disrupting effects in relation to the estrous cycle by BPA using an evidence-based adverse outcome approach^99^. That assessment concluded that BPA decreased estradiol levels in ovarian follicular cells, which subsequently provokes decreased feedback regulation of luteinizing hormone (LH) leading to lengthening of the estrous cycle as well as ovarian cell apoptosis. This clearly shows the added value of using multiple lines of evidence to support an endocrine-mediated effect on female reproductive toxicity. To facilitate this, continued development and validation of robust NAMs for relevant endpoints is required as well as increased availability for OECD-endorsed (quantitative) AOPs and AOP networks^100^.

**Figure 6.**
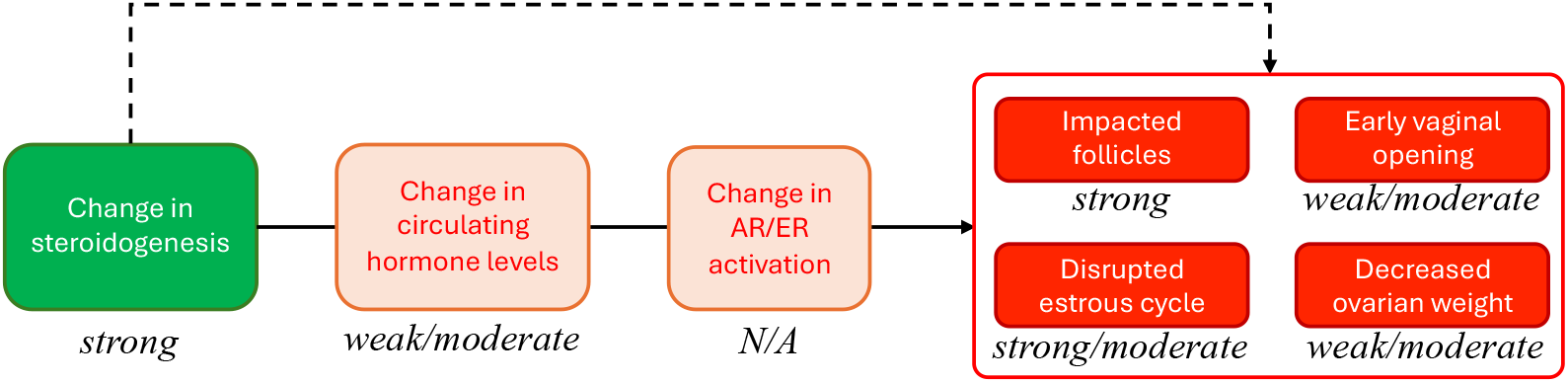
Putative mechanistic pathways leading from disrupted steroidogenesis to female reproductive toxicity. The weight of evidence (weak, moderate, strong) indicated is based on the evidence from present study. N/A not assessed in this study.

## General conclusion

The gold standard to assess steroidogenic disruption is the H295R steroidogenesis assay, as described in the OECD test guideline 456, and currently focuses on changes in T and E2. Studies from our lab and others have shown that culture media obtained from the adrenocorticocarcinoma H295R cell cultures contain a wide range of steroid hormones from the canonical steroidogenic pathway, including various corticosteroids, progestogens, androgens, and estrogens, as well as some backdoor pathway steroid hormones. Results from present study, with 15 blind-coded test compounds, showed that changes in the 15 measured steroid hormone levels in cultures of H295R cells were robust. Comparison of the outcomes with other laboratories, after un-coding of the test compounds, is needed to confirm this. Classification of 14 test compounds did not change when performed based on T/E2 or based on a broader steroidome. One compound was classified as negative for steroidogenic disruption based on T/E2, but showed changes in one backdoor steroid hormone. Considering the current limited knowledge regarding the biological role of the backdoor steroidogenic pathway, interpretation of this finding remains elusive. Expanding the number of hormones as readout in the H295R assay may provide additional mechanistic information to explain potential endocrine-related adverse outcomes. However, it remains to be determined which additional hormones should be included to provide added value to the current H295R assay for regulatory purposes. Additionally, the occurrence of non-monotonic responses and how to use such outcomes for decision-making should be discussed. Finally, more mechanistic information is needed to define clear lines of evidence to link an endocrine effect in an in vitro assay, such as the H295R assay, to female reproductive outcomes in vivo and support regulatory decision-making. This mechanistic information can be obtained from in silico, in vitro or in vivo studies and should preferably be supported by well-developed (quantitative) AOPs and/or AOP networks.

## Acknowledgements

The authors would like to thank the Dutch Ministry of Infrastructure and Water management (IenW) and the independent supervisory committee for providing constructive feedback during the advisory board meetings. We thank Manuel Heinzelmann and Sandra Nijmeijer for the technical assistance and the experimental work performed.

