## Supplemental Tables for "Enhanced H295R steroidogenesis assay and its predictive value for female reproductive toxicity"

Supplementary

**Table S1**. Klimisch categories for reliability of publications using SciRAP assessment

| **Reliability Category** | **Principles** |
| --- | --- |
| **Reliable without restriction** | SciRAP methodological quality score > 80 and all key criteria are “Fulfilled” and there are no deficiencies in the non-key criteria that might affect study reliability. |
| **Reliable with restriction** | SciRAP methodological quality score > 65 and one or several of the key criteria are “Partially Fulfilled” or there are minor deficiencies in the non-key criteria that might affect study reliability. |
| **Not reliable** | SciRAP methodological quality score < 65 or one or several key criteria are “Not Fulfilled” or there are major deficiencies in the non-key criteria that affect study reliability. |
| **Not assignable** | Two or more of the key criteria are “Not Determined”. |

**Table S2**. Principles for categorization in weight of evidence assessment ^101^

| **Category** | **Principle for Categorization** |
| --- | --- |
| **Strong** | Effects were observed in one or more studies judged as reliable without restriction; there may be few conflicting studies, but these do not outweigh the supporting evidence or conflicting results can be explained. |
| **Moderate** | Effects were observed in one or more studies judged as reliable with restriction; there are no conflicting results.  Or effects were observed in one or more studies judged as reliable without restriction or reliable with restriction but with conflicting results, i.e., no or opposite effects were observed in other studies. However, conflicts of results can be explained by differences in study design, for example different exposure periods, doses or animal species or cell models.  There may be few conflicting studies that can not be explained, but these do not outweigh the supporting evidence. |
| **Weak** | Effects were observed in one or more studies judged as reliable without restriction or reliable with restriction but with conflicting results, i.e., no or opposite effects were observed in other studies. Conflicts of results cannot be explained by differences in study design, for example different exposure periods, doses or animal species or cell models.  Or effects were only observed in one or more studies judged as not reliable or not assignable. |
